# Thermal modulation of Zebrafish exploratory statistics reveals constraints on individual behavioral variability

**DOI:** 10.1101/2021.03.17.435787

**Authors:** Guillaume Le Goc, Julie Lafaye, Sophia Karpenko, Volker Bormuth, Raphaël Candelier, Georges Debrégeas

**Affiliations:** Sorbonne Université, CNRS, Institut de Biologie Paris-Seine (IBPS), Laboratoire Jean Perrin (LJP), Paris, France; Université Paris Sciences et Lettres, Paris, France

**Keywords:** behavior, variability, thermokinesis, zebrafish, navigation, locomotion

## Abstract

**Background:** Variability is a hallmark of animal behavior. It contributes to survival by endowing individuals and populations with the capacity to adapt to ever-changing environmental conditions. Intra-individual variability is thought to reflect both endogenous and exogenous modulations of the neural dynamics of the central nervous system. However, how variability is internally regulated and modulated by external cues remains elusive. Here we address this question by analyzing the statistics of spontaneous exploration of freely swimming zebrafish larvae, and by probing how these locomotor patterns are impacted when changing the water temperatures within an ethologically relevant range.

**Results:** We show that, for this simple animal model, five short-term kinematic parameters - interbout interval, turn amplitude, travelled distance, turn probability and orientational flipping rate - together control the long-term exploratory dynamics. We establish that the bath temperature consistently impacts the means of these parameters, but leave their pairwise covariance unchanged. These results indicate that the temperature merely controls the sampling statistics within a well-defined kinematic space delineated by this robust statistical structure. At a given temperature, individual animals explore the behavioral space over a timescale of tens of minutes, suggestive of a slow internal state modulation that could be externally biased through the bath temperature. By combining these various observations into a minimal stochastic model of navigation, we show that this thermal modulation of locomotor kinematics results in a thermophobic behavior, complementing direct gradient-sensing mechanisms.

**Conclusions:** This study establishes the existence of a well-defined locomotor space accessible to zebrafish larvae during spontaneous exploration, and quantifies self-generated modulation of locomotor patterns. Intra-individual variability reflects a slow diffusive-like probing of this space by the animal. The bath temperature in turn restricts the sampling statistics to sub-regions, endowing the animal with basic thermophobicity. This study suggests that in Zebrafish, as well as in other ectothermic animals, ambient temperature could be used to efficiently manipulate internal states in a simple and ethological way.

## Background

Variability, both inter- and intra-individual, is an ubiquitous trait of animal behavior [1]. Intra-individual variability may participate in efficient strategies, as best exemplified by the alternation of exploration and exploitation phases during foraging [2]. It can also endow the animal, or the population, with robustness, *i*.*e*. the ability to rapidly and efficiently cope with changing environmental conditions [3, 4]. The idea, known as bet-hedging, is that a modest loss in fitness associated with phenotypic variability could be balanced by the gain in leniency when facing unexpected and possibly hostile conditions. The origin of inter-individual variability may be attributed to genetic, epigenetic or developmental differences. Intra-individual variability may in turn reflects spontaneous transitions between distinct brain states, *i*.*e*. patterns of persistent neural activity [5, 6]. It may also be the signature of endogenous modulations in the production of neuromodulators [7].

Although the functional significance of variability in animal behavior is now largely recognized [8], the way it is regulated and modulated by external cues, as well as its neuronal substrate remain elusive. To address this question, one not only needs to quantify variability, but also manipulate it in a physiologically relevant manner, in an animal that is accessible to both behavioral and neuronal circuit interrogation. Here we used the zebrafish larva as a model vertebrate as it is uniquely amenable to *in vivo* whole brain functional imaging [9, 10, 11] and to high-throughput behavioral studies [12, 13].

As an ectothermic animal, zebrafish must actively navigate towards regions of its environment that are thermally optimal for its thriving [14], while potentially being exposed to a wide range of temperatures [15]. How fish swim in thermal gradients has been extensively studied [16], and the neuronal circuits underlying this thermotactic process have been identified [17]. Zebrafish larvae integrate thermal signals (change in temperature) over a sub-second time window, and adapt their forthcoming movement accordingly in order to eventually move towards optimal zones.

Here we focus on the exploratory dynamics at various but spatially uniform temperatures. We use a reductive approach, as previously introduced [18], to quantify its spontaneous locomotion using a finite number of short-term kinematic parameters. We then quantify how the bath temperature not only impacts the mean of these parameters, but also their statistical distribution (variability) and pairwise covariance. We further assess the time-scale over which this behavioral variability unfolds at the level of individual animals. From this detailed analysis, we build a numerical model of zebrafish larvae navigation at all temperatures over the physiologically relevant range. Finally, we use this model to demonstrate how this thermal adaptation of spontaneous swimming pattern may complement the thermotactic mechanism, based on direct gradient sensing, in order for the animal to limit its presence in potentially harmful environments.

## Results

### A behavioral assay to record spontaneous navigation at different temperatures

Batches of 10 zebrafish larvae aged 5-7 days were video-monitored at 25 frames/second for periods of 30 minutes as they freely swam in a rectangular 100×45×4.5mm^3^ pool at a constant and uniform temperature (figure 1A, see Methods). For each batch, we successively imposed up to 5 values of temperature (18, 22, 26, 30 and 33°C) in a random order. This thermal range spans the non-lethal conditions for larval zebrafish, and has been shown to be effectively encountered by the animal in its natural habitat [19]. Each 30 min-long recording session was preceded by a 14 min-long period of habituation to allow the animals to reach their steady-state exploratory regime. A total of 10 batches per temperature involving 170 different fish were used.

**Figure 1.**
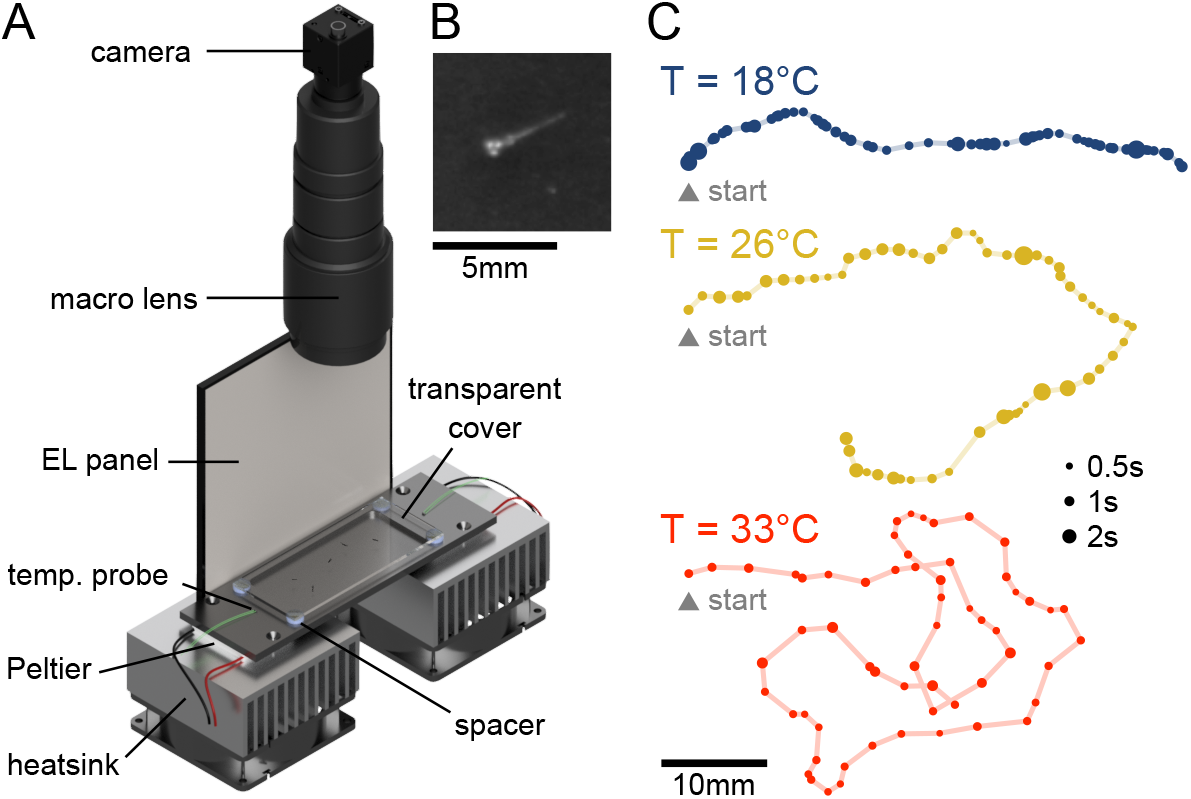
Behavioral assay for the video-monitoring of spontaneous navigation of zebrafish larvae at different temperatures. **A** Sketch view of the setup: Larval zebrafish are freely swimming in a rectangular pool connected to a pair of Peltier modules in a light-tight box. The setup is illuminated with a white electroluminescent (EL) panel and a symmetrically positioned a mirror (not shown). The tank is covered with a transparent slide to limit evaporation. A CMOS camera records images at 25 frames per second. **B** Blow-up of a raw image around a larva. **C** Example trajectories extracted offline from movies recorded at different temperatures. Each dot represents a bout event, with size encoding the time spent at this location.

Larval zebrafish swim in discrete bouts lasting for about 100*ms*, interspersed with ∼ 1 − 2s periods of rest. As we aim to probe how the bath temperature impacts the long-term exploratory process, we focus on the characterization of a few kinematic parameters associated with each bout. We thus ignore the fine structure of the swimming events, such as the amplitude of the tail deflection or beating frequency [20, 21], but examine their resulting heading reorientation and linear displacements. The center of mass coordinates and orientation of each larva in every frame were extracted using FastTrack [22] (see Methods). For each identified swim bout, we computed three scalar parameters (Figure 2A) whose statistics control the fish spatio-temporal exploration [18]: (i) the interbout interval (IBI), *δt*_*n*_, is the idle time following the bout event, (ii) the displacement, *d*_*n*_, is the travelled distance associated with the bout, and (iii) the reorientation angle, *δθ*_*n*_, denotes the change in heading direction.

**Figure 2.**
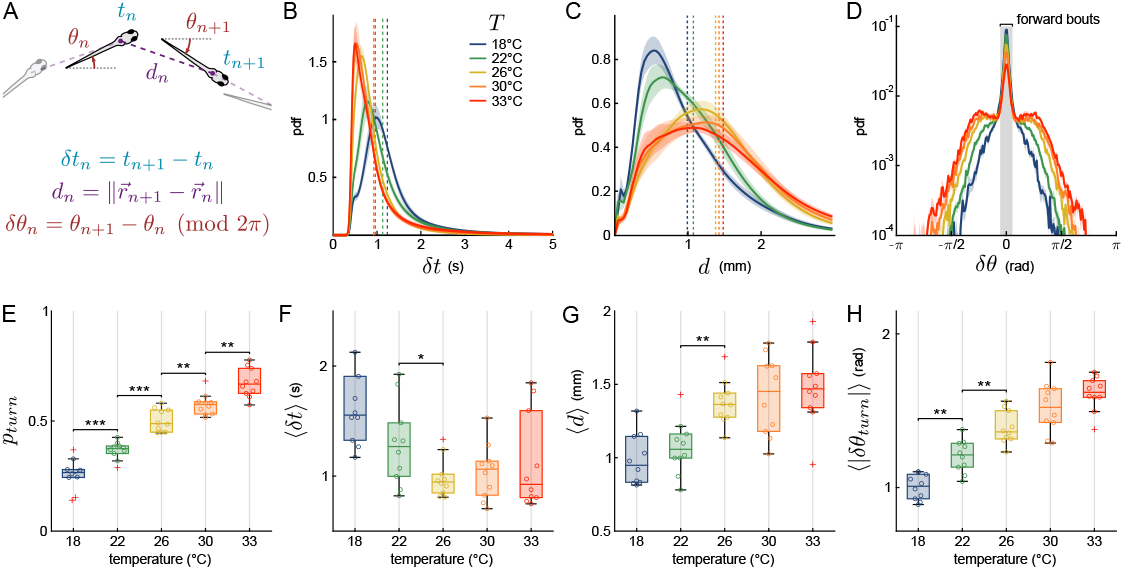
Effects of bath temperature on spontaneous navigation. **A** Sketch defining three kinematic parameters. *δt*_*n*_ is the time elapsed between bout *n* and bout *n* + 1, known as the interbout interval. The displacement *d*_*n*_ is the distance travelled during bout *n* (in mm), while *δθ*_*n*_ represents the reorientation angle. A small value around 0 corresponds essentially to a forward swim, while a large positive value (resp. negative) corresponds to a left (resp. right) turn. **B-D** Per-batch averaged distributions of interbout intervals (**B**), displacements (**C**) and turn angles (**D**) for each tested temperatures. Vertical dotted lines are the means of the distributions, shaded areas are standard errors of the mean (sem). The gray area in **D** marks the forward events versus the turn events. **E-H** Boxplots of selected parameters. Each dot corresponds to a batch of 10 fish, the box spans the 25th to the 75th percentiles, the horizontal line is the median, red crosses are outliers. Significance given only for neighboring boxes (Kruskal-Wallis test, no star : *p >* 0.05, ⋆ : *p* < 0.05, ⋆⋆ : *p <* 0.01, ⋆ ⋆ ⋆ : *p <* 0.001). **E** Fraction of turns, referred to as the turning probability, defined as the ratio of turn bouts over the total number of bouts. **F** Means of the interbout intervals. **G** Means of the displacements. **H** Means of the absolute reorientation amplitude of turning bouts.

Zebrafish larvae, as most animals, tend to adopt a distinct navigational behavior at proximity of boundaries [23]. Tracking was thus performed within the innermost region of the arena, at a minimum distance of 5 mm from the walls. As a result, individual fish were not tracked continuously over the entire recording periods, but along *trajectories* (from one wall to another). In the analysis, we ignored trajectories that last less than 25 seconds. Example trajectories for three temperatures are shown in figure 1C, where each dot indicates the location of a swim bout, while its size reflects the interbout interval. This comparison provides a first qualitative illustration of the effect of temperature on the fish exploration. At low temperatures (18°C), the trajectories are relatively straight, comprising a majority of small discrete forward bouts executed at relatively low frequency. At high temperatures, the trajectories appear much more meandering, with more frequent and ample reorienting maneuvers with longer travelled distances. In the following, we quantify these differences by systematically comparing the statistics of the per-bout kinematic parameters at different temperatures.

### The bath temperature controls the statistical distributions of the kinematic parameters

For each batch and temperature, a probability density function (pdf) was computed for interbout intervals, displacements and turn angles by pooling all bout events. We then computed an average distribution across batches (figure 2B-D, respectively) for the 3 parameters, as well as the temperature-dependence of their mean values (figure 2F-H).

A decrease in the bath temperature from 26°C to 18°C is associated with an increase of the mean IBI (⟨*δt*⟩) from 1 to 1.4s, while the bout frequency remains essentially unchanged at higher temperatures (2B, F). This increase in the mean values is accompanied by a systematic broadening of the statistical distribution. The per-bout displacement exhibits a similar trend (figure 2C). This quantity increases in the range 18-26°C from 1 to 1.5mm, and remains unchanged at higher temperatures (figure 2G).

The turn angle distributions shown in figure 2D reveal the existence of two main bout categories [13, 24, 18]. The central narrow peak corresponds to forward bouts while the wide tail is associated with turning events. We adjusted this distribution as a sum of two empirically chosen functional forms in order to extract the fraction of turning bouts *p*_*turn*_ (see Methods). This quantity steadily increases with the temperature, from 0.3 to 0.8 (figure 2E). This increase in the fraction of turning bouts comes with an increase in their associated reorientation angles *δθ*_*turn*_ as shown in figure 2H.

### The bath temperature controls the persistence time of the orientational state

In a recent study [18], we showed that the orientational dynamics of zebrafish larvae can be described by two independent Markov chains (figure 3A). The first one controls the bout type selection, between forward scoots or turn bouts. This process is essentially memoryless, such that the transition rates are simply set by the ratio between either categories, namely *p*_*turn*_ and 1 − *p*_*turn*_. The second Markov chain controls the orientations of the turning bouts. When a turn bout is executed and if this chain is in the left (right) state, then the animal turns left (right, respectively). This second selection process has been shown to display a persistence over a few bouts: the fish tends to chain turn bouts that are similarly orientated [24, 25, 26], a mechanism whose result is to enhance the angular exploration [18].

**Figure 3.**
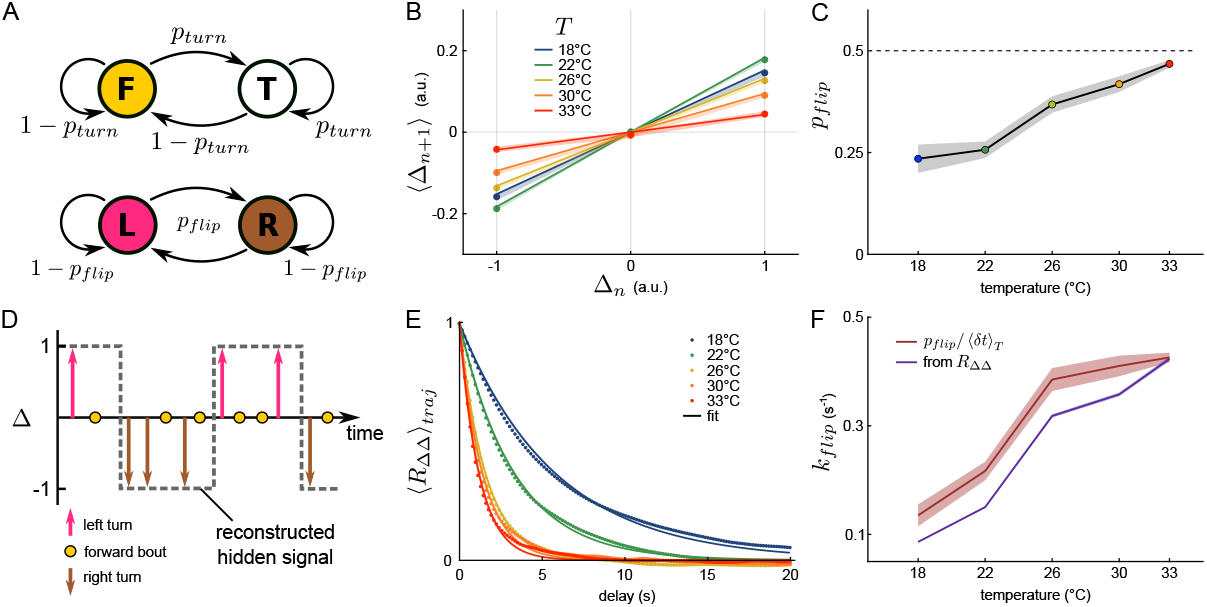
The orientational dynamics is temperature-dependent. **A** Two discrete and independent Markov chains describe the reorientation dynamics. The first one (top) selects the bout type, either turn (T) or forward (F), given the transition rate *p*_*turn*_, while the second one (bottom) determines if the fish is in the left (L) or right (R) state with a transition rate denoted *p*_*flip*_. **B** Mean ternarized reorientation Δ of the next bout, given the current bout reorientation. Shaded area is the sem, solid line is the fit (equation 1). **C** Temperature dependence of *p*_*flip*_. The dashed line at 0.5 indicates a memoryless process. **D** Schematic representing a motion sequence generated by the two discrete Markov chains. The hidden underlying orientational signal that sets the left/right state of the fish is exposed only when the fish performs a turning bout and can be estimated (dashed line) for each trajectory. **E** Trajectory-averaged autocorrelation function of Δ (*R*_ΔΔ_) and associated fit (equation 2). **F** Temperature dependence of *k*_*flip*_, extracted from two methods: *p*_*flip*_ divided by the mean interbout interval associated with each temperature (red, shaded area is the s.e.m.) and from the fit of the autocorrelation function (purple, error bar 95% confidence interval).

Here we examined how this motor-persistence mechanism is impacted by the bath temperature. We estimated the flipping rate *p*_*flip*_ - the probability to switch orientation at each bout - by first binning the turning angles into three categories (denoted Δ) and assigning a discrete value to each of them: right turn (Δ= −1), forward bout (Δ= 0) and left turn (Δ= +1). We then computed the mean discretized angle value (Δ_*n*+1_) at bout *n* + 1 for the three possible values of the previous bout Δ_*n*_, as shown in figure 3B. The slope of the linear fit provides a measurement of *p*_*flip*_ (see Methods and equation 1). This flipping probability increases with temperature from 0.22 at 18°C to 0.45 at 33°C (figure 3C), approaching 0.5. Hence, at high temperatures, the orientational persistence essentially vanishes, *i*.*e*. the probability to trigger a left *vs* a right turn becomes independent of the orientation of the previous bout. This increase of the flipping probability counteracts the concurrent increase in turning probability and turn bout amplitude at high temperature by limiting the tortuosity of the trajectories in these conditions.

This approach yields a typical number of bouts 1*/p*_*flip*_ over which the turning orientation is maintained. A complementary approach consists in characterizing the actual time-persistence (in seconds) of the orientational state [18]. To do so, we assume that the orientation selection is driven by a hidden two-state continuous signal, of which the turn bouts provide a stochastic sampling. We hypothesize that a forward bout is “transparent”, *i*.*e*. it does not interfere with the persistence process, and that the orientational state remains unchanged until a bout in the opposite direction is executed. The procedure for reconstructing the orientational signal is illustrated in figure 3D.

For all trajectories, we computed the autocorrelation function (ACF, *R*_ΔΔ_) of the reconstructed orientational signals, and averaged them for each temperature (figure 3E). The ACF shows a faster decay for higher temperatures, *i*.*e*. the time period over which the animal can maintain its orientational state is larger in colder water. The ACFs could be correctly adjusted with an exponential decay, a functional form that is expected for a simple telegraph process [27]. This suggests that the left/right transition over a time interval *dt* is simply given by *k*_*flip*_*dt*, where *k*_*flip*_ is the transition rate from one state to another. From the exponential fit of the ACFs, we extracted *k*_*flip*_, which we found to increase quasi-linearly with the temperature, as shown in figure 3F (purple line). The rate *k*_*flip*_ is the temporal counterpart of the per-bout flipping rate *p*_*flip*_, the two quantities being linked through the interbout interval. Consistently, we found that *p*_*flip*_/ (*δt*) provides a good approximation of *k*_*flip*_ for all temperatures (figure 3F, red line).

### Navigational kinematic parameters are statistically coupled

In the preceding sections, we showed that the bath temperature impacts in a systematic way the statistical distributions of the five kinematic parameters that control the fish spontaneous navigation, namely the interbout interval (IBI), turn amplitude, travelled distance, turn probability and orientational flipping rate. When examining trajectories recorded at a given temperature, we noticed that they tend to fall in stereotypical categories reminiscent of those most often observed at various temperatures. Some trajectories are tortuous with short IBI, akin to typical hot water trajectories, while other appear to be straighter with less frequent bouts as generally observed in cold water (figure 4A and 1C). This is suggestive of the existence of a finite kinematic repertoire accessible to the animals whose relative occurrence may be controlled by the bath temperature.

**Figure 4.**
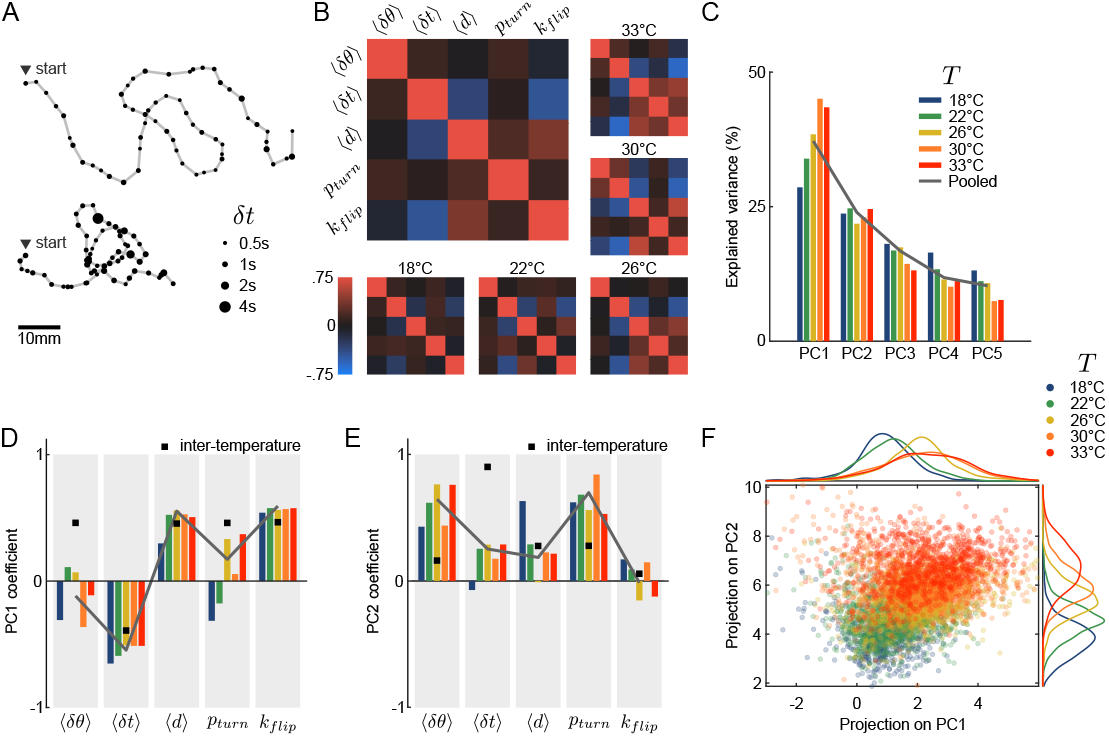
Correlations between parameters are conserved across temperatures. **A** Two qualitatively different trajectories recorded at the same temperature (30°C). **B** Pearson’s correlation matrices of the average reorientation angle *δθ*, interbout interval *δt* and displacement *d*, along with the turning rate *k*_*turn*_ and flipping rate *k*_*flip*_ defined for each trajectory, at different temperatures. Large panel: average over all temperatures. **C** Variance explained by each principal component of a PCA performed on each intra-temperature feature matrix. **D-E** Coefficients of the principal components for intra-temperature matrices (colors), for the inter-temperature averaged matrix (black square) and for the pooled per-temperature array (solid line). **D** First principal component (PC1), **E** second principal component (PC2). **F** All per-trajectory values projected into the principal component space (first two PCs), and their associated marginal distributions for each principal vector.

To test this intuition, we first aimed at establishing the statistical constraints that could set this accessible repertoire. We thus examined the pairwise covariance of the aforementioned kinematic parameters. At short time scale (over one bout), we did not observe any significant correlation between the 3 parameters that can be evaluated on a per-bout basis (IBI, reorientation angle and travelled distance, see Additional file 1: Figure S1A). However, when performing the same analysis on per-trajectory averages, we observed a robust covariance of the parameters. This is illustrated in figure 4B which shows the covariance matrices computed for all data and for each temperature. The IBI appears to be strongly anti-correlated with the forward displacement and the flipping rate. In contrast, besides IBI, all pairs of parameter tend to exhibit positive correlations. Importantly, these statistical features are conserved across the entire temperature range.

### Temperature controls the distribution probability within a well-defined locomotor repertoire

We sought to evaluate how this intra-temperature covariance of the navigational parameters aligned with the inter-temperature covariance. To do so, we used the temperature-averaged parameters to build a 5 temperatures by 5 parameters matrix from which we computed an inter-temperature Pearson correlation matrix (Additional file 1: Figure S1B). The latter displays a comparable structure as the mean intra-temperature correlation matrix 4B: as we have shown in the previous sections, all parameters increase with temperature, and are thus positively correlated, except for the interbout interval which decreases with the temperature and is therefore anti-correlated with the 4 other parameters.

Hence, intrinsic variability and temperature-induced behavioral changes both reflect a concerted modulation of the kinematic parameters along a similar axis. To confirm this claim, we performed a principal component analysis on both the inter-temperature and intra-temperature data. For all temperatures, the first principal component (PC) explains 28 to 45% of the intra-temperature variance (figure 4C), *i*.*e*. significantly more than expected for independent parameters (20%). Due to the small size of the inter-temperature matrix (5 samples), the first PC explains more than 90% of the inter-temperature variance (Additional file 1: Figure S1C). The first PC is conserved across the temperature range (figure 4D, colored bars) and essentially aligned with the inter-temperature PC (black squares). The second PC is similarly conserved across temperatures (figure 4E) yet less clearly aligned with its inter-temperature counterpart.

In order to represent data from various temperatures within the same low-dimensional space, we performed a PCA analysis on the pooled covariance matrix, combining all intra-temperature arrays after standardization (figure 4C-E, solid gray line). Based on the Guttman-Kaiser criterion, we only retained the first two principal components [28] (Additional file 1: Figure S1D). Figure 4F shows the entire dataset projected in this unique 2D PCA space, where the temperature is colorcoded. As the temperature is increased, the accessible locomotor space is shifted towards higher values of both marginal projections, with a concurrent widening of the distribution for the first PC. These observations are thus in line with the view that the trajectories are confined to a manifold defined by the correlation between the various parameters. Each temperature delimits a specific accessible region of this subspace as defined by the PCs projection values.

### Single-fish recordings reveal a slow diffusive-like modulation of the locomotor behavior

The experiments on which these analysis were performed are based on simultaneous recordings of 10 fish for each batch. As we can not track individual fish over the entire session, we can not evaluate to what extent individual animals’ navigational pattern may vary during the course of the assay. To address this specific question, we performed a second series of experiments in which single animals (N = 18) were continuously monitored for 2h at an intermediate bath temperature of 26°C. The same analysis pipeline was implemented. In particular, the recordings were split into successive “trajectories” corresponding to wall-to-wall sequences. We observed that over the course of the assay, the trajectories tended to exhibit strongly distinct features as illustrated in figure 5A, reflecting a significant intra-individual behavioral variability.

**Figure 5.**
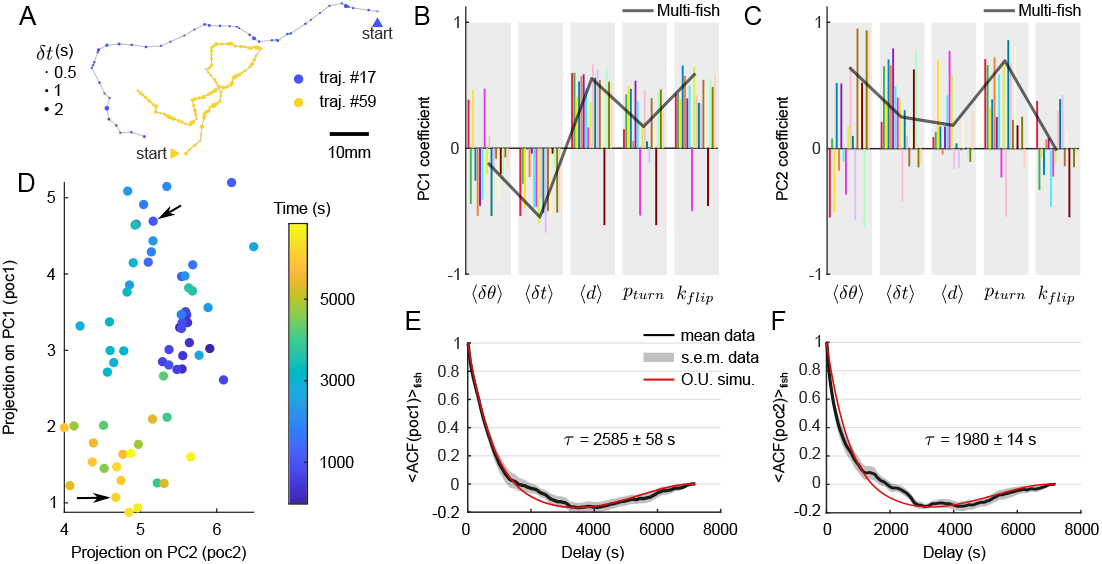
Diffusive-like exploration of the behavioral manifold for individual fish. **A** Two qualitatively different trajectories from the same fish at the same temperature (26°C), recorded at 1h interval. **B-C** Coefficients of the two first principal components for 18 different fish (one color corresponds to one fish). The solid line is the PC coefficients computed from the multi-fish experiments as shown in figure 4E-F. **D** Time-evolution of the projections in the 2D PCA space from an example fish. One dot corresponds to one trajectory whose parameters are projected on the multi-fish PC space. Color encodes the time at which the trajectory starts. Arrows show trajectories represented in **A** with the same color. **E-F** Autocorrelation function of the projections on (**E**) PC1 and (**F**) PC2, averaged across fish. Gray area is the standard error of the mean. Red line is the autocorrelation function of a simulated Ornstein–Uhlenbeck process whose bias parameter (1*/τ*) is fitted to the data.

For each individual, we similarly computed a feature matrix containing, for all successive trajectories, the mean interbout interval, reorientation angle of turn events, displacement, turning probability and flipping rate. We then performed a PCA on each array. Both the explained variance (Additional file 2: Figure S2A) and the PCA coefficients (figure 5B-C) were unchanged with respect to the multi-fish analysis (5B-C, gray line). This indicates that the covariance structure in the locomotion pattern is similar at the intra and inter-individual level.

We thus used the multi-fish PC space defined in the preceding section to represent the single-fish data. The result for an example fish is shown in figure 5D where the successive trajectories are indicated as dots in this two-dimensional PC space. This representation reveals a slow diffusive-like exploration of the locomotor space over the course of the experiment, with a progressive transition from one type of trajectory (e.g. long displacements, frequent bouts, frequent turns) to another (e.g. short displacements, longer inter-bout intervals and fewer turns).

To quantify the time-scale of this itinerant exploration within the locomotor space, we computed the autocorrelation function (ACF) of the projections on the two first PCA components (5E-F, black line). These curves could be captured by an Ornstein–Uhlenbeck (O.U.) process, which describes the dynamics of a random walker within a quadratic energy basin [29, 30], see Methods). The latter allows one to bound the stochastic exploration within a finite region of the locomotor space. From the fit, we extracted the times needed for the dynamical system to reach its stationary regime: *τ* = 2585±58 s for PC1, *τ* = 1980±14 s for PC2 (mean ± s.e.m.). These values clearly demonstrate that the modulation of the exploratory behavior in individual animals takes place over time scales that are orders of magnitude longer than the interbout interval.

This series of experiments allowed us to further assess the relative contribution of the intra- and inter-individual components in the observed behavioral variability. As the assay is longer (2h) than the time needed to reach the stationary regime (∼ 2000*s*), each recording provides an estimate of the intra-individual variability. The latter was quantified in the PC space as the variance of the PC projections across the entire duration of the assay, averaged over the various individuals. We then separately computed the variance of the PC projections, pooling the data of all animals (Additional file 2: Figure S2D, green). The latter quantity thus encompasses both inter- and intra-individual variability. This analysis led to the conclusion that a dominant fraction of the variance (68% on PC1, 53% on PC2) can be explained by the intra-individual variability.

### Simulations of spontaneous navigation at various temperatures reveal basic thermophobic behavior without direct gradient-sensing mechanism

Having thoroughly characterized the statistical structure of the kinematic parameters and their thermal modulation, we sought to build a minimal stochastic model of the fish navigation in order to generate synthetic trajectories at different temperatures. Each kinematic parameter defines a random variable whose mean is set by the temperature and whose statistical distribution accounts for both the inter-trajectory variability and the per-bout stochasticity. The dual nature of the variability was mathematically recapitulated by expressing each of the 5 kinematic variables as a product of two stochastic, temperature-independent variables: one accounting for the trajectory-to-trajectory modulation (within a range controlled by the bath temperature, Additional file 3: Figure S3B-E, *Y* column), and the other reflecting the remaining short-term variability (bout-to-bout, Additional file 3: Figure S3B-E, *ϵ* column, see Methods). For the former, we used the copula method to reproduce the observed covariance of the per-trajectory means of the various parameters.

This approach allowed us to generate various trajectories at different temperatures, as illustrated in figure 6A. These trajectories are qualitatively similar to those typically observed at the corresponding temperatures (see figure 1C for a comparison). To quantify how this stochastic model captures the exploratory behavior, we computed the mean square displacement (MSD, figure 6B) and the mean square reorientation (MSR, figure 6C) on both the real (dots) and numerical data (solid lines). Overall, the exploratory dynamics appear to be correctly reproduced by the numerical model. Importantly, the inter-trajectory variability is also, by construction, correctly reproduced by this minimal model.

**Figure 6.**
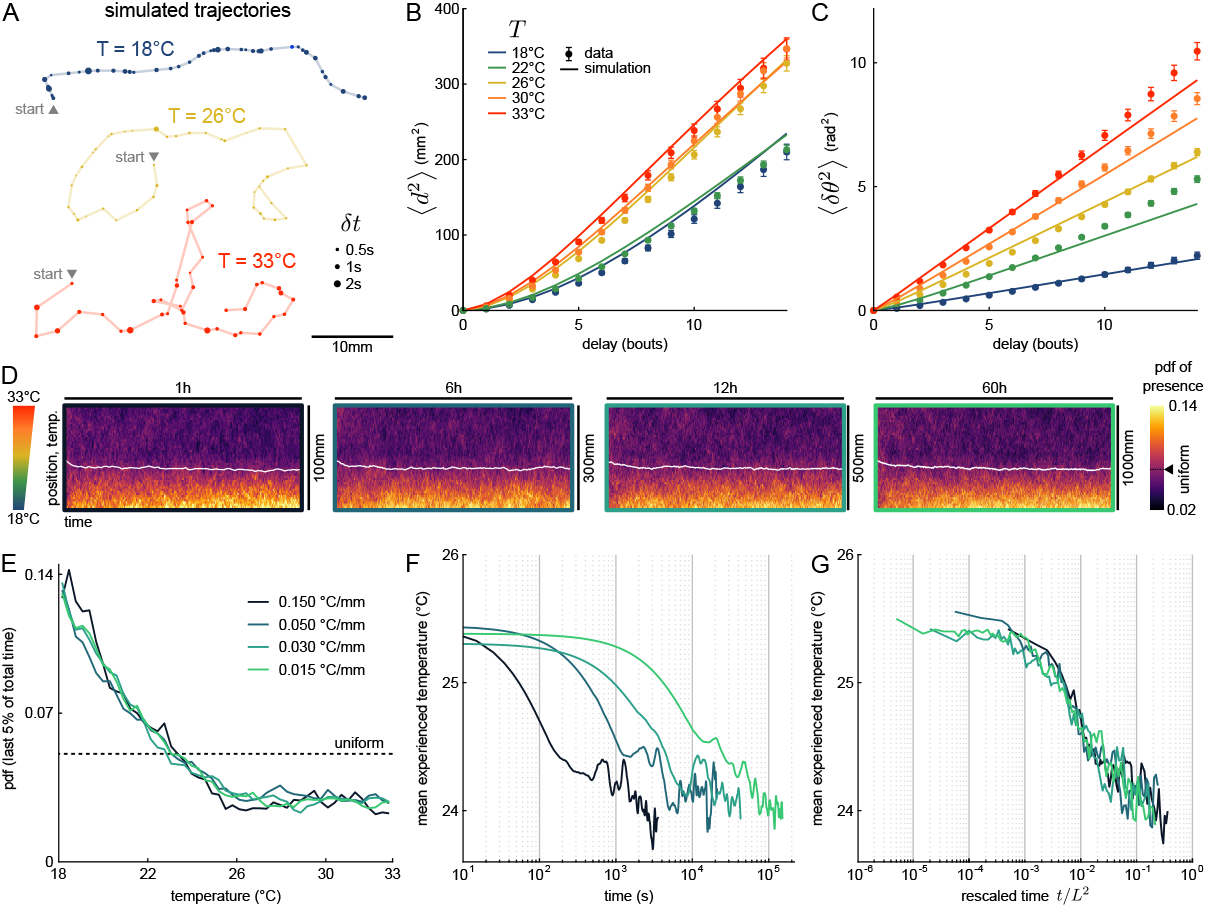
Simulations indicate that zebrafish does not need gradient information to perform negative thermotaxis. **A** Example trajectories generated with a simulation based on rescaled multivariate distributions (see Methods). **B** Mean square displacement, from data (dots) and simulation (line). **C** Mean square reorientation, from data (dots) and simulation (line). **D** Distributions of presence of simulated fish through time, for four strengths of temperature gradient. The white curve is the average position over time. The expected value for a uniform distribution is highlighted on the colormap. **E** Steady-state distribution of presence as a function of temperature. The dashed line is the expected value for an uniform distribution. **F** Temporal evolution of the average position over time (only the first 75 bins are shown for readability). **G** Distribution mean as a function of the time rescaled by the squared pool length.

This model was used to probe how the temperature dependence of the navigational kinematics may participate in driving the animal along thermal gradients. We first experimentally quantified how zebrafish larvae responded to a linear thermal gradient spanning our temperature range (18°C-33°C), by focusing on the steady-state occupation distribution. We found that the larvae favor regions where the temperature is comprised between 23°C and 29°C (Additional file 4: Figure S4), *i*.*e*. they tend to avoid both extreme (hot and cold) regions. The underlying sensory-motor mechanism is bound to involve both the effect of the temperature on the fish navigation pattern (thermokinesis) and a direct (immediate) response to perceived temperature changes (thermotaxis) [16, 14]. Our model allows us to assess the relative contribution of the kinesis process. In order to do so, we implemented a simulation in which a virtual fish navigates in a rectangular pool (*L* × 45mm) in which we imposed a linear thermal gradient along the horizontal *x*-axis spanning the 18°C-33°C range. We simulated trajectories of numerical swimmers by continuously updating their exploratory statistics according to the local bath temperature. These changes are entirely controlled by the temperature-dependence of the 5 kinematic parameters, which we linearly interpolated across the thermal gradient. Four gradient strengths were emulated by changing the length *L* of the pool (*L* = 0.1, 0.3, 0.5, 1*m*).

The time evolution of the position distribution along the gradient are shown as heatmaps in figure 6D. They reveal a global drift of the population towards the low temperature region for all values of the thermal gradient (figure 6E). In all conditions, the distributions were found to converge towards a unique steady-state profile after a finite time. The probability of presence in the steady-state regime displays a quasi-linear decay from 18 to 26°C, and remains uniform at higher temperature. The thermokinesis process thus endows the animal with a basic thermophobic behavior, even for minute gradients - orders of magnitude smaller than those imposed in thermotactic assays. In contrast, the avoidance of cold regions seen in experiments (Additional file 4: Figure S4, see Methods) is absent in our simulations, and must therefore reflect a direct gradient-sensing mechanism.

The dynamic of this thermophobic behavior in the simulations appears to depend on the imposed gradient, as illustrated in figure 6F, which shows the mean experienced temperature across the population as a function of time for the three gradients. All the curves display a similar decay associated with a global drift towards the cold region, until a similar plateau value is reached, albeit with different time-scales. Due to the diffusive nature of the fish spatial exploration, the settling time is expected to scale with the square of the pool length. Consistently, the four dynamic evolution are found to fall on a unique curve when plotted as a function of *t/L*^2^ (figure 6G). The associated settling times range from 10 minutes for the largest gradient up to ∼ 14 hours for the smallest one.

## Discussion

Animal behaviors unfold as trajectories in a high dimensional space of motor actions. To make behavior mathematically tractable, one needs to unveil statistical rules that couple the different components of the behavior and organize them across time-scales. This dimensionality reduction approach is a pre-requisite to further distinguish between deterministic and stochastic components of the behavior and concurrently discover the underlying neural mechanisms [31, 32]. Leveraging novel techniques for high-throughput behavioral monitoring and automatic classifications has allowed to elucidate the statistical structure organizing self-generated behaviors in numerous species, such as *C. elegans* [33], *Drosophila* [34, 35], zebrafish [21, 36], or mice [37].

With its bout-based navigation, zebrafish larva offers a relatively simple model for such an endeavour. It has been shown that as few as 13 different swim bout types are sufficient to capture the entirety of its behavioral repertoire [21]. Here we focus on spontaneous exploration in the absence of time-varying sensory cues. For this particular behavior, Marques et al. showed that only 4 different bout types effectively contribute to the navigation. In the present work, we further restricted the discretization to only two categories (forward and turn bouts), and extracted 5 kinematic variables, ignoring fine differences in tail bout execution. We showed that the knowledge of these 5 variables statistics nevertheless suffices to fully characterize the long-term exploratory process. Indeed, synthetic trajectories generated by stochastic sampling from the statistical distributions extracted from the data accurately reproduce the experimentally observed angular and translational dynamics.

Using this reductionist approach, we were able to demonstrate that the variability in the fish exploratory dynamics originates from two separate mechanisms, acting on distinct time-scales. Over a few bouts, the fish displacement is akin to a random walk in which multiple stochastic processes set the successive values of two discrete (bout type and turn bout orientation) and three continuous (Inter-bout-interval, linear and angular displacements) variables that together define its instantaneous in-plane velocity. These processes are statistically constrained by mean transition rates and amplitude probability distributions that can be considered invariant at the scale of individual trajectories (*i*.*e*. over tens of bouts). These parameters however vary significantly over long time scales: their time modulation takes place over hundreds to thousands of bouts, indicative of a clear time-separation between the two different processes. Importantly, although we did not observe any significant correlation in the instantaneous locomotor variables, the slow modulation of the kinematic parameters exhibits robust covariance, and is thus constrained within a well-defined kinematic manifold.

The present study allowed us to quantify how the water temperature modulates the locomotor statistics of zebrafish larvae. Rather than evoking distinct locomotor patterns, temperature controls the relative occupancy within this subspace: changing the temperature consistently impacts the mean value of the kinematic parameters but leaves their covariance structure unchanged. Temperature thus essentially sets the accessible range of exploratory trajectories within a well-defined continuum of possible locomotor behaviors.

At the circuit level, it is tempting to interpret these observations by considering the brain as a dynamical system exhibiting multiple metastable patterns of activity (brain states) whose relative stability and transition rates define a particular energy landscape [38]. In this view, the short-time dynamics that select the successive bout properties correspond to a stochastic itinerant exploration of this neuronal landscape. The latter is essentially invariant over minutes but is slowly reshaped via endogenous processes or through temperature changes, leading to a gradual modification of the short-term statistics.

Slow modulation of locomotor characteristics in zebrafish larvae have been reported in two recent studies [2, 36]. In [2], the authors identified two discrete states, associated with exploration and exploitation during foraging, with typical persistent times of order of minutes. In [36], progressive changes in locomotor statistics were associated with decaying hunger state, as the initially starved animal progressively reached satiety. In contrast with these two studies, the modulation in locomotor kinematics that we observed is continuous and does not reflect spatial heterogeneities in the environment (e.g. local presence of preys) or explicit changes in internal states such as satiety. With respect to hunger state, the use of temperature may offer a practical way to externally drive the internal state to a stationary point in an ethologically relevant way.

The neuronal basis of this internal state modulation process remains to be elucidated. The circuits regulating specific locomotor features, such as the bout frequency [39] or orientational persistence [25, 26] have been identified. However, the fact that the various kinematic parameters display concerted endogenous modulations points towards a global drive. Temperature is known to impact cellular and synaptic mechanisms [40] in such a way that an increase in temperature tends to speed up neuronal oscillatory processes [41, 42]. This may explain the concurrent decrease in the persistent times associated with the orientational persistence and interbout intervals. The thermal modulation of the angular and linear amplitude of the bouts may in turn reflect a temperature dependence of the muscular efficiency rather than neuronal processes [43]. Another possibility is that the temperature drives the activity level of neuromodulatory centers which may also exhibit slow endogeneous modulations. This neuromodulation release would then globally impact the spontaneous dynamics of various premotor centers yielding the observed change in locomotor patterns. The serotonergic neurons of the dorsal raphe constitute an attractive candidate for such a mechanism as their activation has been shown in numerous instances to drive a persistent change in behavior in zebrafish [44, 5, 2], as well as in mice [45].

Our study yields a minimal numerical model of zebrafish locomotion at different temperatures. This model allowed us to probe *in silico* how the thermal modulation of the exploratory dynamics may contribute to the thermotaxis behavior, thus complementing direct gradient-sensing mechanisms [17]. Our simulations indicate that this thermokinesis process endows the animal with the capacity to efficiently avoid hotter regions, but cannot explain the observed avoidance of cold water. As thermal gradient sensing operates within a time window of 400ms [16], it may be ineffective in conditions where the lengthscale of thermal gradients is much larger than the typical distance travelled per bout. In such conditions, this complementary mechanism may be strategically relevant as it allows the animal to navigate away from potentially noxious regions.

## Conclusions

This study establishes the temperature as an effective and practical external parameter to explore behavior variability in vertebrates. Our analysis provides simple latent variables, namely the two first PCA projections, that can be used to efficiently track the animal’s behavioral state. Changes in behavioral states are generally induced through complex protocols, involving a perturbation of a sensorimotor loop, or through abrupt changes in sensory conditions [46]. In such approaches, the change is discrete and generally transient as the animal eventually adapts to the new conditions. In contrast, temperature offers a way to drive a robust, continuous and chronic shift in behavior that can be easily implemented while performing large-scale brain monitoring. Various behavioral states are thought to reflect different levels of attention or arousal, which in turn impact the responses to sensory stimulation. Beyond its utility for studying how a given neuronal circuit may give rise to distinct dynamics, thermal perturbation could also be leveraged to investigate how internal states may enhance or inhibit sensory responses.

## Methods

### Animals maintenance

Experiments were performed with wild type *Danio rerio* AB, aged 5 to 7 days post-fertilization (dpf). Larvae were reared in Petri dishes containing embryo medium (E3), at 28°C, with a 14/10 hours cycle of light/dark and were fed with nursery powder GM75 everyday from 6dpf. Experiments were done during daytime, in E3. They were approved by Le Comité d’Éthique pour l’Expérimentation Animale Charles Darwin C2EA-05 (02601.01).

### Experimental setup

The container consists in a rectangular pool (100×45×2.5 mm^3^) made of copper whose surface was protected by a biocompatible heat-resistant black layer (Rust-Oleum). It is stuck on two 78W Peltier modules (Adaptive) with thermal tape (3M). A transparent, 2mm-thick PMMA cover is placed over the pool with 2mm spacers to minimize water evaporation, leaving a water thickness of 4.5mm. The temperature is measured at both ends of the pool with thermocouples type T (Omega). The two left/right error signals (*T*_*target*_ − *T*_*measured*_) are used within two independent PID loops implemented on an Arduino Uno board (Arduino) whose coefficients have been optimized manually. Each PID regulates the PWM frequency sent to a H-bridge driving the power sent to the two Peltier modules. A graphical user interface (GUI) written in C++ using the Qt framework is used to monitor the measured temperatures and to impose the target temperatures on both ends. Due to its high thermal diffusivity, the copper piece quickly reaches a uniform temperature and acts as a thermostat for the water. After about 4 minutes, the temperature of the water in the center of the pool has reached the set temperature (±0.2°C), which then remains constant over time. The GUI monitors the bath temperatures while grabbing frames from a CMOS camera (FLIR Chameleon3 CM3-U3-13Y3M-CS) coupled wih a macrolens (Navitar) at 25 frames per second. The whole apparatus is placed in a light-tight box, illuminated with a homogeneous white light emitted by a LED panel (Moritex) placed on the side; a mirror placed at the other side limits significant phototactic bias in the small direction of the pool. All codes mentioned above are available on Github [47] under a GNU GPLv3 licence. Blueprints of the box and pool as well as electronic designs are available upon request.

### Innocuity of the black-painted copper pool

Zebrafish larvae are sensitive to minute concentration of chemicals. To check the innocuity of the container, ten zebrafish larvae were left overnight inside the setup. All survived and were swimming actively. We further checked whether the black layer may releases chemical compounds that could impact the animals navigational dynamics. We prepared a stock of E3 heated for 1h at 45°C in the experimental setup. 3 batches of 10 larvae were then sequentially recorded for 30 minutes in E3 (control), in a Petri dish, then placed for 40 minutes in the heated E3 and recorded again for 30 minutes (treated). Both experiments were performed at 28°C. We then computed the mean interbout interval, mean displacement, mean reorientation amplitude for each batch and the cumulative mean square displacement. Additional file 5: Figure S5A shows that none of these quantities display any significant change for larvae recorded in the heated water with respect to the control. Copper being known to alter lateral line neuromasts [48], we further tested the absence of copper ions by incubating 10 larvae in the heated E3 for 2 hours before marking their lateral line neuromasts with DiAsp according to the protocol detailed in [48]. All larvae displayed intact lateral line neuromasts.

### Experimental protocols

The pool is filled with E3. A temperature is randomly drawn from 18, 22, 26, 30, 33°C and set with the GUI. After 10 minutes, a batch of 10 zebrafish larvae is introduced in the pool. After 10 minutes of habituation, the fish kinematics are monitored for 1800s (half an hour). In order to confirm the full habituation of the animal to the new conditions, we checked that the fish navigation is indeed time-invariant during the recording period. We evenly split each recording into 3 time windows and computed the statistics of the various kinematic parameters (mean bout frequency, displacement and reorientation amplitude) for these 3 periods, The distributions within each time-window are not significantly different (*p >* 0.1, two-sample Kolmogorov-Smirnov test) as shown in Additional file 5: Figure S5B. Fish remain in the pool while we randomly draw a new non-tested temperature. After 20 minutes (temperature regulation and habituation), a new recording of 1800s is performed. The five temperatures are not systematically tested on all batches, but for each temperature, 10 different batches of 10 fish are used. To check whether the testing order may impact the kinematic of swimming for a given temperature, we separated the assays in two groups depending on whether the previous recording was performed at a lower (Δ*T >* 0) versus higher (Δ*T <* 0) temperature. We then computed the mean interbout intervals, mean displacement and mean reorientation amplitude normalized by their temperature mean. The comparison of the two distributions is shown in Additional file 5: Figure S5C. For all parameters, the statistical difference is non-significant, indicating that the testing order has no major impact on the navigational statistics. In total, the experiments involved 17 different batches. The sample size was not statistically determined beforehand.

For single-fish experiments, the same protocol is used except that a single fish was placed in the pool. The recordings last for 2h and only T=26°C is tested.

For thermal gradient experiments (Additional file 4: Figure S4), 10 larvae are used during 45 minutes. The first 5 minutes are recorded with a uniform temperature of 22°C, then a linear gradient is imposed during 40 minutes, from 18°C to 33°C. The gradient direction (*i*.*e*. which side is set to either 18°C or 33°C) is chosen randomly. 10 different batches are tested. The distribution of presence along the gradient is computed over the last 2 minutes (5% of the gradient duration) such as to allow enough time for the animals to reach a steady-state.

### Tracking and basic analysis

Larvae were tracked offline using the open-source FastTrack software [22]. It generates a text file containing the position of each fish’s center of mass and body angle across frames until they leave the defined ROI. Kinematic analyses were performed using MATLAB (R2020a, Mathworks). Bouts are detected when the instantaneous speed is greater than two times the overall variance of the speed. Putative bouts are then filtered on a distance criterion (bouts with a linear displacement - measured in a time window of ±0.5*s* centered on the bout onset - less than 0.3mm or greater than 18mm are rejected) and on a temporal criterion (bouts occurring within 0.4s after a bout are rejected). Bout timing is defined as 80ms before the velocity peak. Detection performance was checked manually on randomly selected sequences. From positions, time and body angles before and after a bout event, we computed displacements, interbout intervals, and turn angles associated with each bout. Data are split into trajectories, from one edge of the ROI (set at 5mm from the walls) to another. Only trajectories that last at least 25 seconds, with at least 10 bouts, with 3 bout types (left turn, right turn and forward scoot) are kept for further analysis. Trajectories last on average 67s (median 47s, 95th percentile 178s) and contain on average 60 bouts (median 44 bouts, 95th percentile 154 bouts). The number of trajectories and the number of bouts retained for further analysis are given for each temperature in table 1. The effect of the chosen cutoffs (minimum trajectory duration and minimum interbout interval) used for trajectory and bout selection is tested in Additional file 6: Figure S6. This control demonstrates that changing these cutoffs has no significant impact on our results. All MATLAB routines are available on Gitlab [49] under the GNU GPLv3 licence.

**Table 1.** Number of trajectories, number of bouts and mean trajectory duration kept for analysis per temperature.

|  | Temperature (°C) |  |  |  |  |
| --- | --- | --- | --- | --- | --- |
|  | 18 | 22 | 26 | 30 | 33 |
| # traj. | 617 | 1405 | 1533 | 1308 | 1079 |
| # bouts | 34931 | 96365 | 89955 | 77203 | 48836 |
| duration (s) | 100 | 82 | 56 | 60 | 50 |

### Bout classification

To discriminate whether a bout falls in the forward or the turning categories, we fitted the one-sided (absolute value) reorientation angles distributions with the sum of a zero-mean Gaussian distribution and a gamma distribution. The Gaussian corresponds to the part of the distribution close to zero, while the gamma function aims at describing the distribution of high angle reorientations. We manually set the Gaussian width and the scale parameter of the gamma function based on the observed distributions. We fitted the shape parameter for each temperature, ensuring that the slope at high angles in logarithmic scale is well reproduced. Then, we defined a fixed threshold for the angles to be considered as a turn or a forward bout. This threshold is the angle at which the two distributions cross, invariably found around 10° (10.25 ± 0.23°, mean ± sd). This value of 10° (0.17rad) was used to classify bouts throughout this work.

### Displacement correction

We noticed that the displacement corresponding to a turn event was systematically larger than the displacement associated to a forward event. This is due to the fine structure of a turning bout: first, the fish performs a small reorienting bout, then it scoots forward [20]. Since we do not look at this fine structure, the overall displacement during a turn bout is geometrically overestimated and would bias temperature-to-temperature comparison. We computed the ratio between displacements during turns and the ones during forward swims, and found a factor of 1.6 ±0.1, regardless of the temperature. Therefore, in all analyses presented in this work, all displacements corresponding to a turn event were corrected by a factor 1/1.6 = 0.625.

### Statistical methods

Probability density functions (pdf) were computed with a kernel density estimation through the built-in Matlab function `ksdensity`, with a bandwidth of 0.1 for interbout intervals and displacements and 0.5 for turn angles. For the distributions of figure 2, a pdf was computed for each batch and the mean and standard error of the mean are computed. For rescaled curves (Additional file 3: Figure S3), data from all experiments were pooled to compute the temperature-average quantity 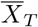 and rescaled values. Boxplots were made with the built-in Matlab function `boxchart`, using as input data the means of the respective quantities for trial (one dot corresponds to a batch of 10 fish). For simulations of navigation, averages over temperature were computed by pooling all bout events from all experiments corresponding to this particular temperature. *p*_*turn*_ and *p*_*flip*_ values were estimated for each trajectory and then averaged. Error bars for those temperature averages and for the pdf shown in Additional file 3: Figure S3 were all computed using bootstrapping with 1000 boots to get the 95% confidence interval through the built-in `bootci` function. Errors were propagated for the ratio of *p*_*flip*_ and ⟨*δt*⟩_*T*_ in figure 3F.

### Reorientation dynamics

The two Markov chains model has been described in details in a previous study [18]. We first binned the reorientation angles *δθ* into a ternarized reorientation Δ, with values −1 (right turn *R*), 0 (forward bout *F*) and +1 (left turn *L*). To extract *p*_*flip*_, we analytically derived the mean reorientation Δ_*n*+1_ given the previous reorientation Δ_*n*_. There are 9 combinations of bouts {*n*; *n* + 1}: {*L*; *L*}, {*L*; *R*}, {*L*; *F*}, {*F*; *L*}, {*F*; *R*}, {*F*; *F*}, {*R*; *L*}, {*R*; *R*}, {*R*; *F*}. All combinations involving a forward bout yield 0. Remain combinations with two turns in the same direction and two turns in the opposite direction. For a turn in direction *L* (resp. *R*), the associated probability corresponds to the case where a flip occurred (*i*.*e*. the previous bout was in direction *R*, resp. *L*) and the case where no flip occurred (*i*.*e*. the previous bout was in direction *L*, resp. *R*). Noting 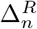 and 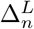 the turns in the right and left direction at bout *n*, the mean reorientation given the direction of the previous bout reads :

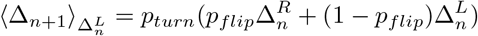

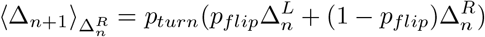

These equations can be summed up as:

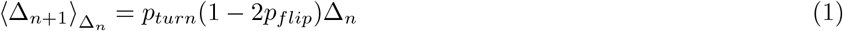

This is the fit used in figure 3B.

A random telegraph signal is a binary stochastic process with constant transition probability per unit of time. In the case where both states are equiprobable, the two transition rates (here noted *k*_*flip*_) are equal. For such processes, the time spent in one or the other state (left or right) is exponentially distributed [27] and the autocorrelation function for a zero-mean signal reads :

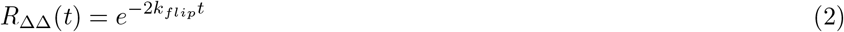

This is the fit used in figure 3E.

Mean square displacement (MSD) ⟨*d*^2^⟩ and mean square reorientation (MSR) ⟨*δθ*^2^⟩ were computed using the MATLAB package `msdanalyzer` [50]. All (*x, y*) and *δθ* sequences are pooled by temperature for both data and simulations, the MSD and MSR were computed for each sequence and we show in figure 6B-C the ensemble average for each temperature with the standard error of the mean.

### Principal components analysis

The “features matrices” were built for each temperature. They include, for each trajectory, mean interbout intervals, turn probability, flip rate (estimated as *p*_*flip*_*/* ⟨*δt*⟩, *p*_*flip*_ being extracted as explained above, for each trajectory), mean reorientation angle of turning events and mean displacements. Each set was standardized (centered and normalized by its standard deviation) before being processed by the single value decomposition (SVD) algorithm through the built-in `pca` function. Those 5 intra-temperature standardized arrays are then concatenated to form the so-called pooled matrix, that is in turn used to find a common space through PCA. For projection, each set was normalized by the standard deviation of all the pooled data (regardless of temperature) and not centered for comparison purposes. The aforementionned common space was also used to project data from single-fish experiments.

### Numerical Ornstein–Uhlenbeck process

The single-fish experiments contains an average of 48 ± 16 trajectories (mean ± s.d.) with a median duration of 42s and a value of the 90th percentile of 122s. Each trajectory yields one point in the PC space. We then linearly interpolated the PCA projection values in order to produce a discrete signal defined over the same time vector across the total duration of the assay (7200s), sampled every second. For each fish, on both PC, we computed the autocorrelation function (Additional file 2: Figure S2B-C) before averaging them (figure 5E-F, black line is the mean, shade is the s.e.m.). The autocorrelation function of a OU process reads [30]:

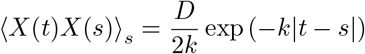

where *D* is the diffusion coefficient and *k* is the bias term. However, this expression is only valid in the limit where the recording time is much larger than the relaxation time 1*/k* of the process. When the recording duration is of order of the auto-correlation decay time, the computed autocorrelation function exhibits a negative overshoot, which reflects an incorrect estimate of the long-time mean of the signal. This issue is common to stochastic signals whose mean is unknown. In order to fit the experimental autocorrelation and extract the relaxation time *τ* = 1*/k*, we used simulated dynamics over similar time-windows. Numerical simulations of the Ornstein–Uhlenbeck (O.U.) process were sequentially implemented using the following equation [51] :

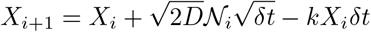

with *δt* the time interval chosen for the simulation (units *s*) and N is a random number drawn from a normal distribution.

To determine *τ*, we generated 500 realisations of the O.U. process with *D* set to 1 and *τ* set to values in a given range. For each realisation, we computed the autocorrelation function (ACF) and averaged them across realisations. We then computed the residual sum of square (RSS) and chose the minimum one to select the best parameter *τ*. After manually narrowing down the best range for *τ* (PC1: 2000s to 3000s, 1000 values; PC2 : 1900 to 2100s, 1000 values), we repeated the previous process 20 times to get 20 “best *τ*” and we report the mean ± s.e.m. in the text and figure.

It should be noticed that the auto-correlation decay times extracted from this analysis are 2585 and 1980s, for PC1 and PC2 respectively. These time-scales are thus one order of magnitude larger than the typical trajectory duration. This clear time-scale separation a posteriori validates the per-trajectory discretization used in our analysis.

### Numerical simulations of trajectories

Trajectories were simulated using the framework described in Additional file 3: Figure S3, based on the hypothesis that (1) spatio-temporal dynamics can be reproduced solely from five parameters, (2) per-bout values of interbout intervals (*δt*), displacements (*d*) and turn angles (*δθ*) are drawn from a distribution that can be decomposed as 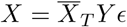, (3) the per-trajectory values of turning probability (*p*_*turn*_) and flipping probability (*p*_*flip*_) are drawn from a distribution that can be decomposed as 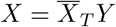 and (4) the trajectory-averaged parameters are correlated. Note that for the simulations we use *p*_*flip*_ rather than flipping rate for simplicity in the code implementation.

#### 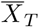, the temperature average

All per-bout values of *δt, d*, reorientation angle of turn events (*δθ*_*t*_) and reorientation angles of forward events (*δθ*_*f*_) are pooled by temperature and the mean is computed. A *p*_*turn*_ and a *p*_*flip*_ is estimated for each trajectory, pooled by temperature and averaged (Additional file 3: Figure S3B-E, left column).

#### Y, the trajectory means variability

For each trajectory, a mean value is computed for *δt, d* and *δθ*_*t/f*_ while *p*_*turn*_ and *p*_*flip*_ are extracted. They are then rescaled by the corresponding temperature average value computed above. For each temperature, a cumulative density function (cdf) is computed. They are then averaged across temperatures to get a single *Y* cdf for each parameters (pdf shown in Additional file 3: Figure S3B-E, middle column).

#### ϵ, the per-bout variability

Similarly, for each trajectory we rescale values of *δt, d* and *δθ*_*t/f*_ by their corresponding trajectory mean. Then, all events are pooled by temperature and a cdf is computed. Finally, we will use the mean cdf, resulting in a single *ϵ* cdf for per-bout parameters. *p*_*turn*_ and *p*_*flip*_ are defined for a trajectory, hence they do not have bout to bout variability (pdf shown in Additional file 3: Figure S3B-D, right column).

#### Correlations of means

We compute the Pearson’s correlation matrix of the trajectories’ parameters (trajectory means and probabilities), for each temperature. The coefficients are then averaged to get a single correlations matrix ⟨*R*_*traj*_⟩_*T*_. *Algorithm*. After choosing a number *n* of fish (trajectories), we generate multivariate distributions (copulas) with the MATLAB built-in `mvnrnd` function, with the mean ⟨*R*_*traj*_⟩_*T*_ correlations matrix as input. It produces 5 marginal sets of *n* gaussian random numbers, correlated with one another. We then get the corresponding normal cdf, which is in turn used to sample the corresponding *Y* cdfs, inversing the latter. Finally, those samples are multiplied by the corresponding temperature average 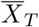. A bout is generated by sampling a displacement and a turning angle, along with a interbout interval during which the virtual fish stands still, from the generic cdf of *ϵ*. Those values are multiplied by the trajectory means drawn earlier, and the new position (*x, y*) is computed. The next bout is generated, and so on. For the gradient simulations, the same strategy is used, at the notable difference that the temperature averages are determined dynamically given the position of the agent along the temperature gradient. We used reflective boundary conditions. We checked the consistency between parameters distributions from the data and from the simulations, as well as correlations between trajectory means.

## Abbreviations

IBI: interbout interval

## Acknowledgements

We thank the IBPS fish facility staff for the fish maintenance, in particular Stéphane Tronche and Alex Bois. We are grateful to Carounagarane Dore for his contribution to the design of the experimental setup, Geoffrey Migault for engineering expertise and Raphaël Voituriez for fruitful discussion on the modeling aspects.

## Availability of data and materials

All data generated or analysed during this study are included in the Dryad repository https://doi.org/10.5061/dryad.3r2280ggw, which also includes MATLAB scripts used to generate figures shown in this article.

## Ethics approval and consent to participate

Not applicable.

## Consent for publication

Not applicable.

## Authors’ contributions

G.L.G., V.B., R.C. and G.D. conceived the project. R.C. designed the setup. G.L.G. and J.L. performed the experiments. G.L.G., S.K. and G.D. analyzed the data. All authors contributed to the manuscript. All authors read and approved the final manuscript.

## Additional Files

Additional file 1 — Figure S1. *Additional file 1*.*pdf*

**Figure S1.**
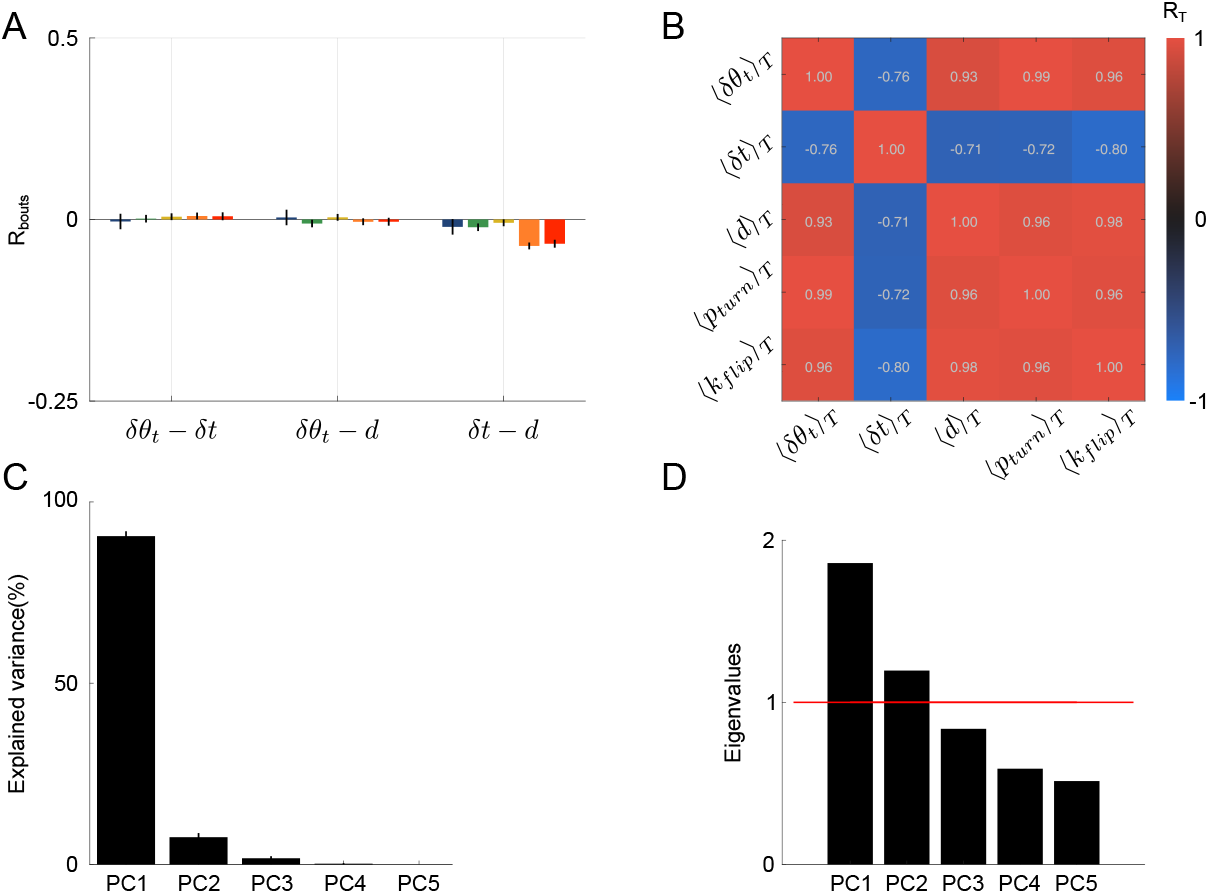
Correlations between parameters. **A** Pearson’s correlation coefficients between per-bout parameters, reorientation angles of turn bouts, interbout interval and displacement. **B** Pearson’s correlation matrix between temperature-averaged parameters. **C** Variance explained by the principal components of the inter-temperature matrix. **D** Eigenvalues of the pooled intra-temperature matrix. The red line highlights the Kaiser-Guttman criterion.

Additional file 2 — Figure S2. *Additional file 2*.*pdf*

**Figure S2.**
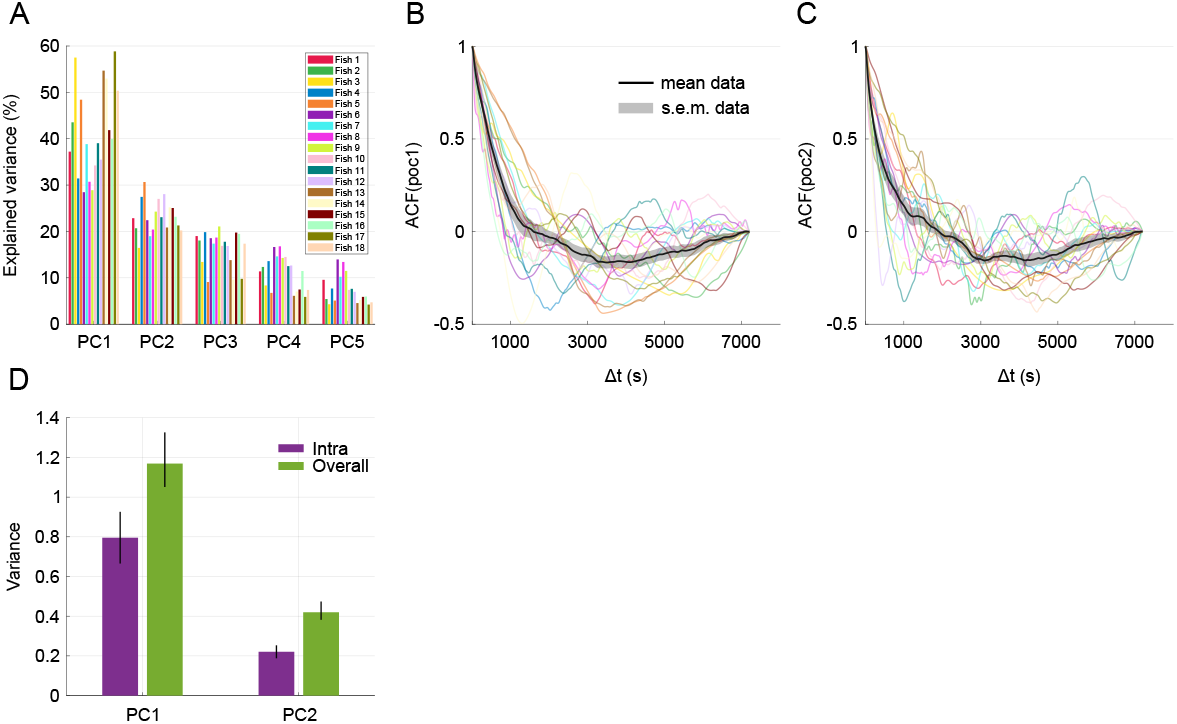
PCA in single-fish experiments. **A** Variance explained by the five principal components for each single-fish. **B-C** Autocorrelation function of the projection on PC1 (B) and PC2 (C) from each fish in single-fish experiments. The color code is the same as in A, black line and shaded area is the mean and s.e.m. across fish. **D** Mean variance of projections across time (intra, purple) and overall variance of projections (green). Error bars for intra is the s.e.m. and error bars for overall is 95% confidence intervals after bootstrapping (n=1000 boots).

Additional file 3 — Figure S3. *Additional file 3*.*pdf*

**Figure S3.**
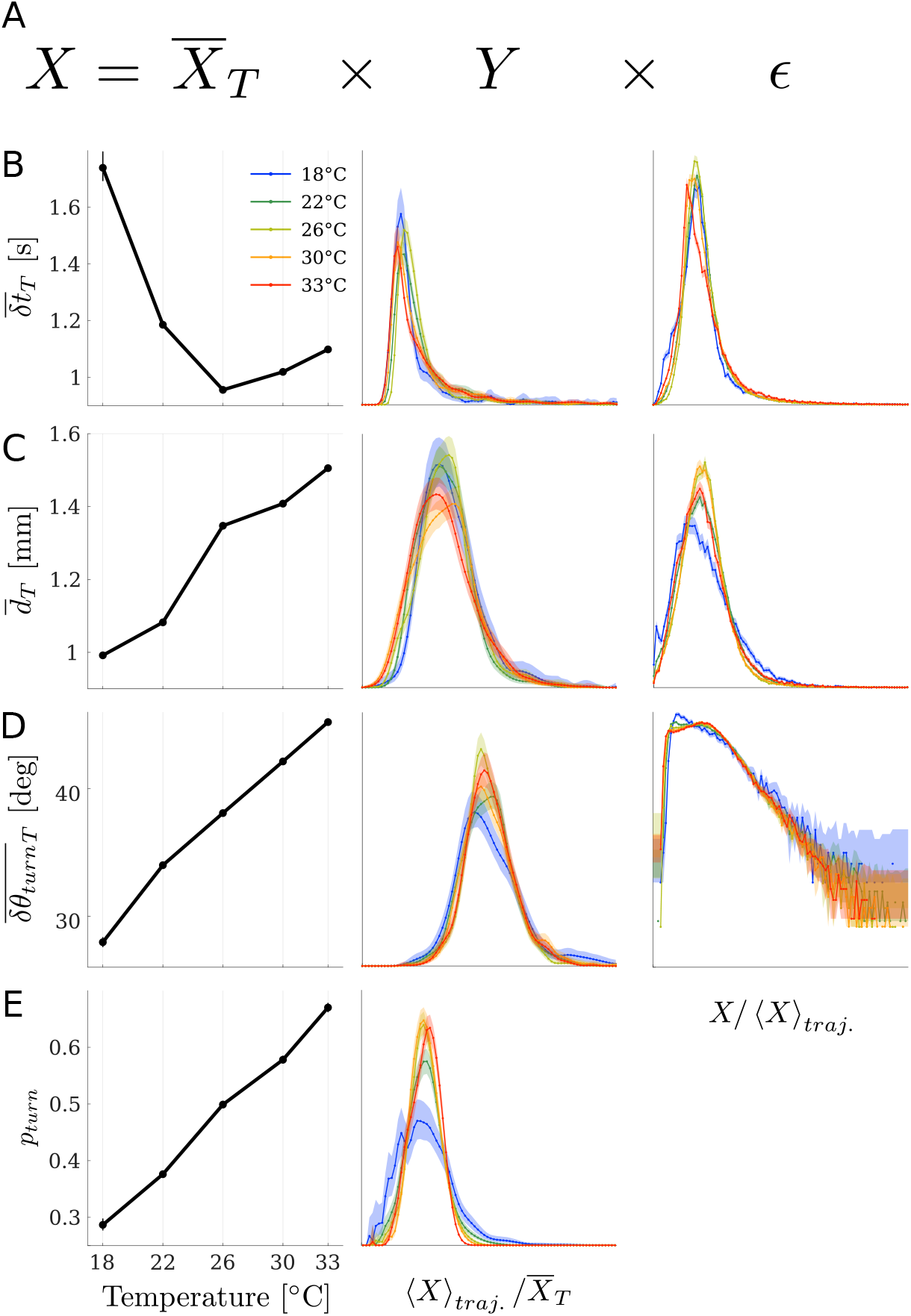
Temperature-independant rescaling of parameters. **A** Equation describing parameter X distribution. **B-E** Left to right, temperature-averaged value, trajectory-averaged rescaled by temperature averaged-value and per-bout value rescaled by the trajectory average, for **B** interbout intervals, **C** displacements, **D** reorientation angle of turn events, **E** turning probability.

Additional file 4 — Figure S4. *Additional file 4*.*pdf*

**Figure S4.**
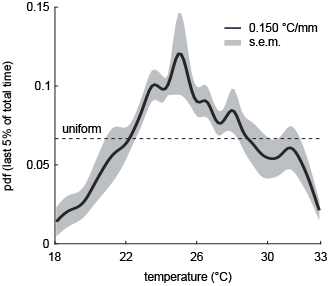
Fish position distributions along a linear thermal gradient. Presence probability density function of 10 batches of 10 larvae experiencing a thermal gradient from 18°C to 33°C. Solid line is the mean across batches, shaded area is the s.e.m. Dashed line is the expected value for a uniform distribution.

Additional file 5 — Figure S5. *Additional file 5*.*pdf*

**Figure S5.**
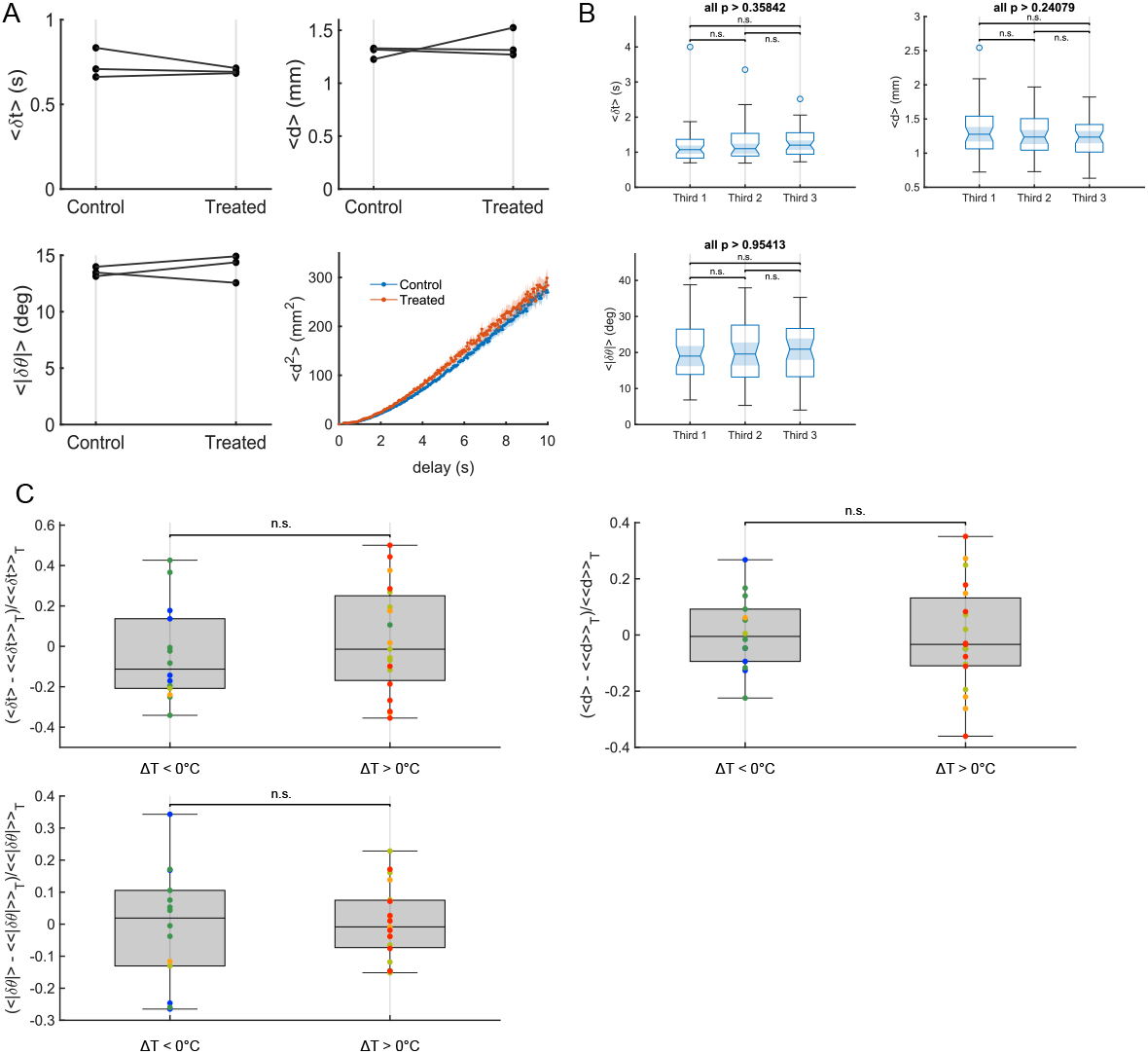
Controls. **A** Example statistics of three batches successively recorded in E3 (control) and in E3 previously heated at 45°C for one hour in the experimental setup. (Left to right, up to bottom) Mean interbout interval, mean displacement, amplitude of turn bouts reorientation angle and mean square reorientation. **B** Example statistics from all experiments pooled together, dividing time into three windows of 10 minutes. (Left to right, up to bottom) Mean interbout interval, mean displacement, amplitude of turn bouts reorientation angle. **C** Example statistics at a given temperature (current T) as a function of the previously tested temperature. One dot is the mean parameter value for one experiment, color encoded current temperatures, lines are the mean for each previous temperature. (Left to right, up to bottom) Mean interbout interval, mean displacement, amplitude of turn bouts reorientation angle.

Additional file 6 — Figure S6. *Additional file 6*.*pdf*

**Figure S6.**
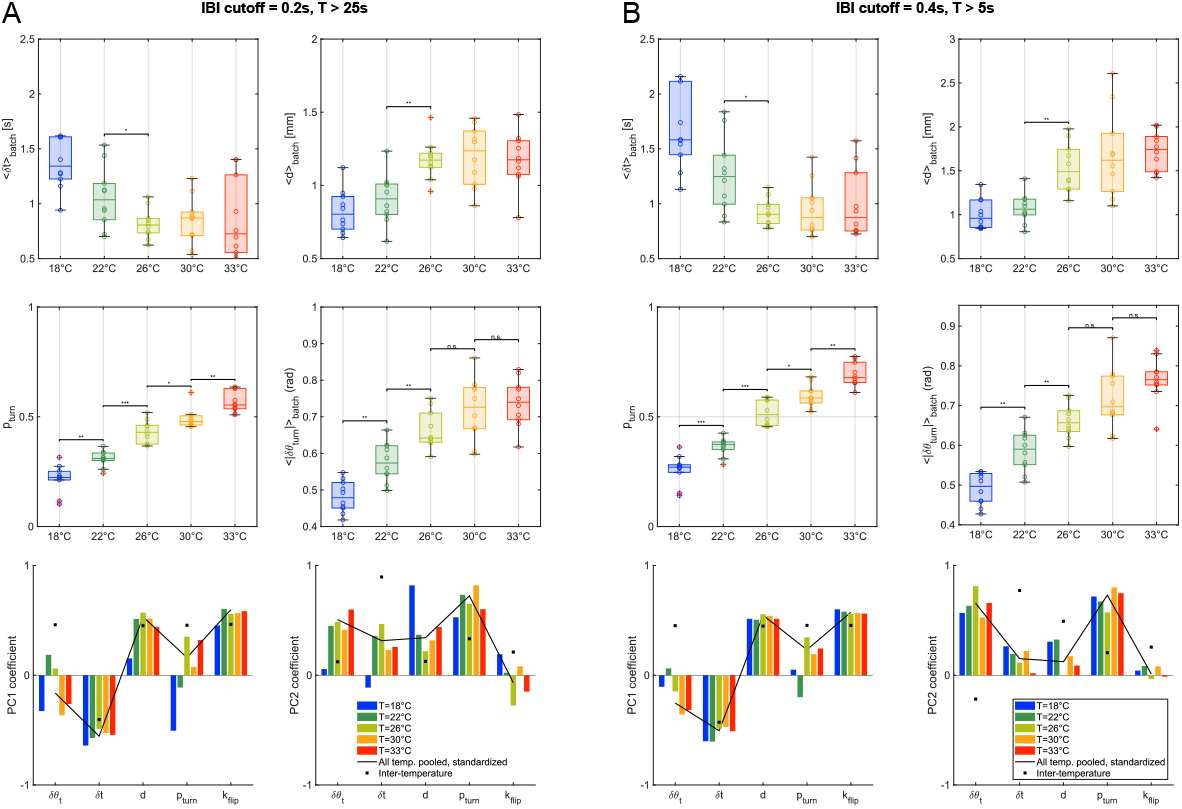
Effect of cut-offs in trajectory selection. **A-B** Example statistics when changing cutoffs in trajectory selection. (Left to right, up to bottom) Mean interbout interval, mean displacement, fraction of turns, amplitude of turn bouts reorientation angle, first principal component coefficients, second principal component coefficients. **A** With a minimum time between two consecutive bout of 200ms, trajectory must last at least 25 seconds. **B** With a minimum time between two consecutive bout of 400ms, trajectory must last at least 5 seconds.

